# Peptide inhibitors targeting FOXO4-p53 interactions and inducing senescent cancer cell-specific apoptosis

**DOI:** 10.1101/2024.09.04.610228

**Authors:** Donghoon Kang, Yeji Lim, Dabin Ahn, Jaeseok Lee, Chin-Ju Park

## Abstract

Cellular senescence, marked by irreversible cell cycle arrest and the secretion of proinflammatory factors, contributes to aging and cancer recurrence. Chemotherapy can also induce senescence; senescent cells often resist apoptosis and promote tumor recurrence. The interaction between FOXO4 and p53 sustains cell survival. Herein, we used biophysical techniques and conducted cellular experiments to develop a peptide inhibitor targeting this interaction to eliminate senescent cancer cells. We identified key regions in the p53 transactivation domain (TAD) involved in FOXO4 binding and designed an optimized peptide inhibitor (CPP-CAND) with improved cell permeability. CPP-CAND showed high selectivity and potency in inducing apoptosis in senescent cells by disrupting FOXO4-p53 foci and activating caspase pathways. It is effective against senescent cancer cells induced by doxorubicin and cisplatin, highlighting its potential as a senolytic drug. Thus, CPP-CAND is a promising therapeutic candidate with improved selectivity, efficacy, and cost-effectiveness.

## Introduction

Organisms lose their physical functions and reproductive capacity after adulthood, ultimately leading to death. This is due to the accumulation of DNA damage and various stress factors inside and outside the cell^1^. Irreversible and severe DNA damage in normal cells is typically eliminated via the apoptosis pathway^2^. Reactive oxygen species, ionizing radiation, and certain chemicals can cause single- and double-strand breaks in DNA^3,4^. Base excision repair and nonhomologous end joining are involved in repairing these damages ^3,4^. If the DNA damage is too severe or there is a defect in the DNA repair pathway, DNA repair will ultimately fail^5-7^. Failure of DNA repair induces apoptosis through p53 activation, mitochondrial pathway induction, and persistent DNA damage response (DDR) signaling^8-10^. Simultaneously, some cells undergo cell cycle arrest and become senescent owing to the inhibition of cyclin-dependent kinases mediated by p21 activation through the DDR pathway^11^.

Senescent cells secrete the senescence-associated secretory phenotype (SASP), which consists of inflammatory cytokines and chemokine growth factors, into the surrounding tissues. This creates an inflammatory environment and promotes cell cycle arrest, inducing senescence in neighboring tissues^12,13^. Cellular senescence is a phenomenon observed not only in normal cells but also in cancer cells^14-16^. Cancer cell senescence is primarily caused by oncogene-induced senescence (OIS) or therapy-induced senescence (TIS). OIS is characterized by genetic alterations that promote cellular aging and inhibit cancer cell proliferation. A prominent example of OIS is the H-Ras^G12V^ mutation, which triggers senescence by inducing the DDR mediated by ROS^17^. On the other hand, TIS can be induced by anticancer drugs and radiotherapy. These treatments eliminate cancer cells by inducing DNA damage, a common anticancer strategy^18,19^. Insufficient levels of treatment do not cause apoptosis but may trigger senescence in cancer cells^20-22^. Chemotherapy activates the DDR pathway. DDR pathway activation phosphorylates and activates p53, which subsequently induces the expression of p21^23^. This cascade of molecular events culminates in the establishment of cellular senescence. For instance, doxorubicin, a topoisomerase inhibitor, induces DNA damage in cancer cells by interfering with topoisomerases and unwinding supercoiled DNA^24^. However, low concentrations of doxorubicin do not induce apoptosis in cancer cells, but rather induce cellular senescence^25,26^. Cancer cell senescence induced by anticancer treatments contributes significantly to cancer recurrence by enhancing cancer invasiveness and stemness^14,27^. Therefore, an effective chemotherapy regimen should aim to eliminate both cancerous and senescent cancer cells to minimize the risk of recurrence.

Several drug candidates have been developed to eliminate senescent cells. Senescent cells are generally characterized by their resistance to apoptosis; therefore, drugs targeting EPHB1, PI3K/AKT, and Bcl-2/Bcl-xL, which are involved in the apoptosis resistance pathway, have been developed^28^. The combination of dasatinib and quercetin (DQ) has completed phase 1 clinical trials for Alzheimer’s disease, and navitoclax is currently in phase 3 clinical trials for myelofibrosis^29,30^. However, drugs such as navitoclax and DQ can cause side effects, such as thrombocytopenia, neutropenia, and pulmonary hypertension, as they also affect normal cells^31,32^. Recently, the interaction between forkhead box O4 (FOXO4) and p53 has received considerable attention as a target for senescent cells. FOXO4 is a forkhead box O transcription factor that mediates various signaling pathways to maintain cellular homeostasis^33^. FOXO4 is localized within the promyelocytic leukemia body in senescent cells and interacts with p53, inhibiting its nuclear exclusion^34^. Consequently, this interaction suppresses the apoptosis of senescent cells induced by p53 in the mitochondria^35^. Additionally, the FOXO4-p53 interaction contributes to the maintenance of senescent cells by activating the transcription of their common target gene, p21cip1, which suppresses apoptosis^36^. Therefore, FOXO4 is a promising target for the development of senolytics. Although FOXO4 is present in a small fraction of normal cells, its expression is significantly increased in senescent cells^34^. Taking advantage of these properties, several protein-protein interaction inhibitors have been developed to eliminate senescent cells. For instance, FOXO4-D-Retro-Inverso (DRI) induces the selective apoptosis of senescent cells by inhibiting the FOXO4-p53 interaction^34^. However, It has a high synthesis cost because of its long length (46 a.a) and the need for D-type amino acids. Furthermore, FOXO4-DRI targets the tumor suppressor p53, a crucial protein, which may lead to various side effects in clinical trials^37^. Another drug, ES2, targets FOXO4 conserved region 3 (CR3) but has a significantly lower selectivity than FOXO4-DRI^25^.

In this study, we designed a peptide inhibitor based on the p53 transactivation domain (TAD) sequence and investigated its binding interface at the atomic level using NMR spectroscopy. Using heteronuclear nuclear Overhauser effect (hetNOE) experiments, we determined the essential regions of the p53 TAD for binding to the FOXO4 forkhead domain (FHD). Next, we employed a fluorescence polarization assay (FPA) to pinpoint the sequences crucial for binding among the specified regions, subsequently improving the cell applicability of the peptide sequence. Finally, we validated that the peptide inhibitor exhibited robust selectivity for senescent cells and efficiently disrupted FOXO4-p53 foci formation despite its short length and L-type amino acid composition. Our study provides a novel peptide drug designed to eliminate senescent cancer cells based on the structural information of the FOXO4-p53 interaction.

## Results

### The p53 TAD2 region plays a crucial role in the interaction with FOXO4 FHD

Interactions between FOXO4 and p53 in the nucleus have been reported to promote cellular senescence^34^. This interaction is mediated by dual binding, with FHD mainly binding to the p53 TAD and FOXO4 CR3 binding to the p53 DNA-binding domain (DBD)^38^ (Figure. 1).

**Figure 1.**
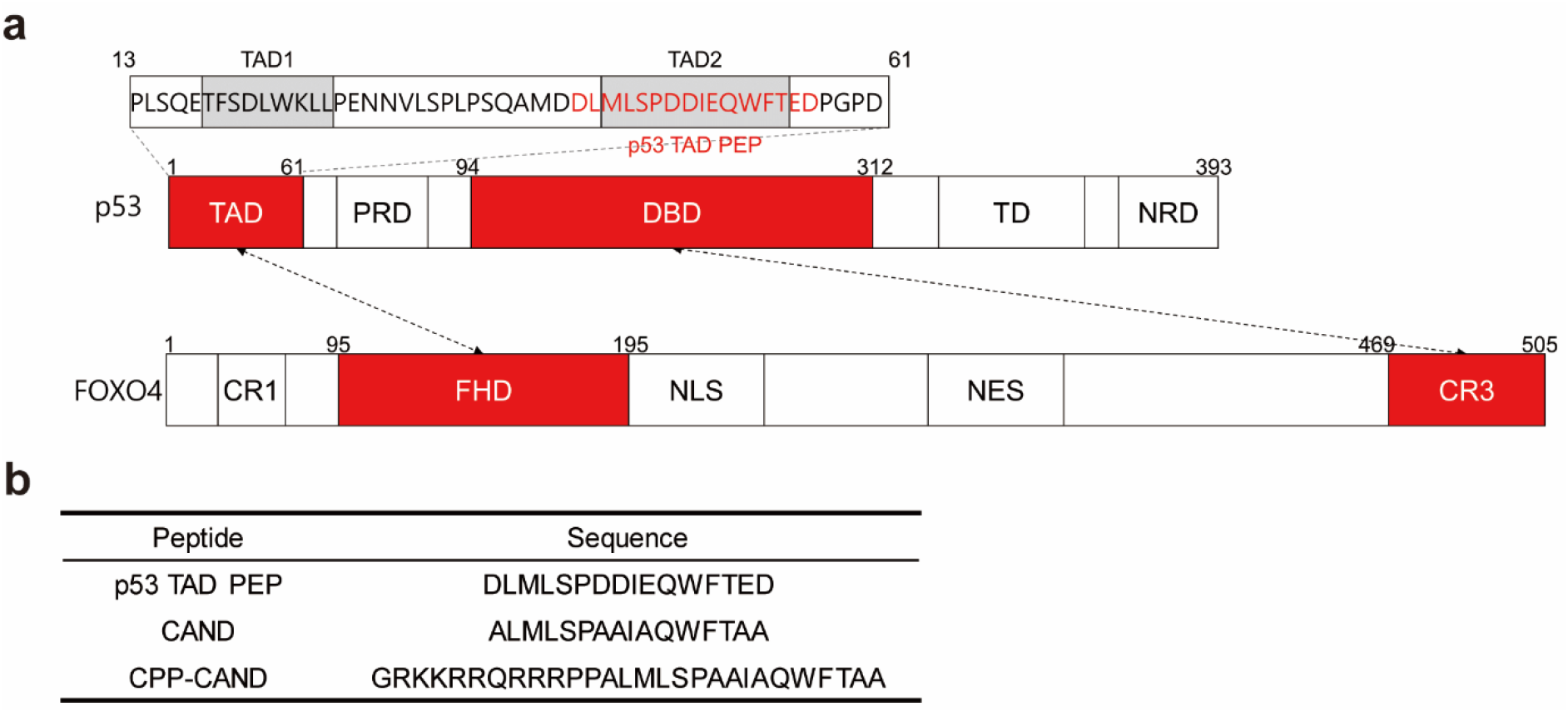
The domain structure of FOXO4 and p53 proteins. (a) Primary sequence of p53 TAD. The arrows connect the domains that are known to interact. The peptide sequence of p53 TAD PEP is indicated in red. TAD, transactivation domain; PRD, proline-rich domain; DBD, DNA-binding domain; TD, tetramerization domain; NRD, negative regulatory domain; CR1 and CR3, transactivation domains; FHD, forkhead domain; NLS, nuclear localization sequence; NES, nuclear export sequence. (b) Sequences of p53 TAD PEP, CAND, and CPP-CAND peptides. The CPP-CAND sequence was designed by introducing HIV-TAT sequences to CAND.

Paramagnetic resonance enhancement (PRE) experiments were conducted to investigate the interaction between the FOXO4 FHD and p53 TAD. We used ^15^N-labeled FOXO4 FHD and p53 TAD with S-(1-oxyl-2,2,5,5-tetramethyl-2,5-dihydro-1H-pyrrol-3-yl)methyl methanesulfonothioate (MTSL)-labeled Cys46 for the PRE experiments. MTSL caused peak broadening near the labeled residues, allowing us to identify which region of the FOXO4 FHD was close to the p53 TAD. We found a notable decrease (less than 0.2) in the peak intensity at the N-terminus and helices 3 and 4 of the FOXO4 FHD (Figure. 2a, b). It was noteworthy that approximately half of the residues exhibiting a substantial decrease in intensity were hydrophobic amino acids (Figure. 2b). This result is consistent with previous investigations of the interaction between FOXO4 FHD and p53 TAD using chemical shift perturbation (CSP)^38^. This suggests the significance of the hydrophobic nature of the interface in the FOXO4 FHD and p53 TAD interaction.

**Figure 2.**
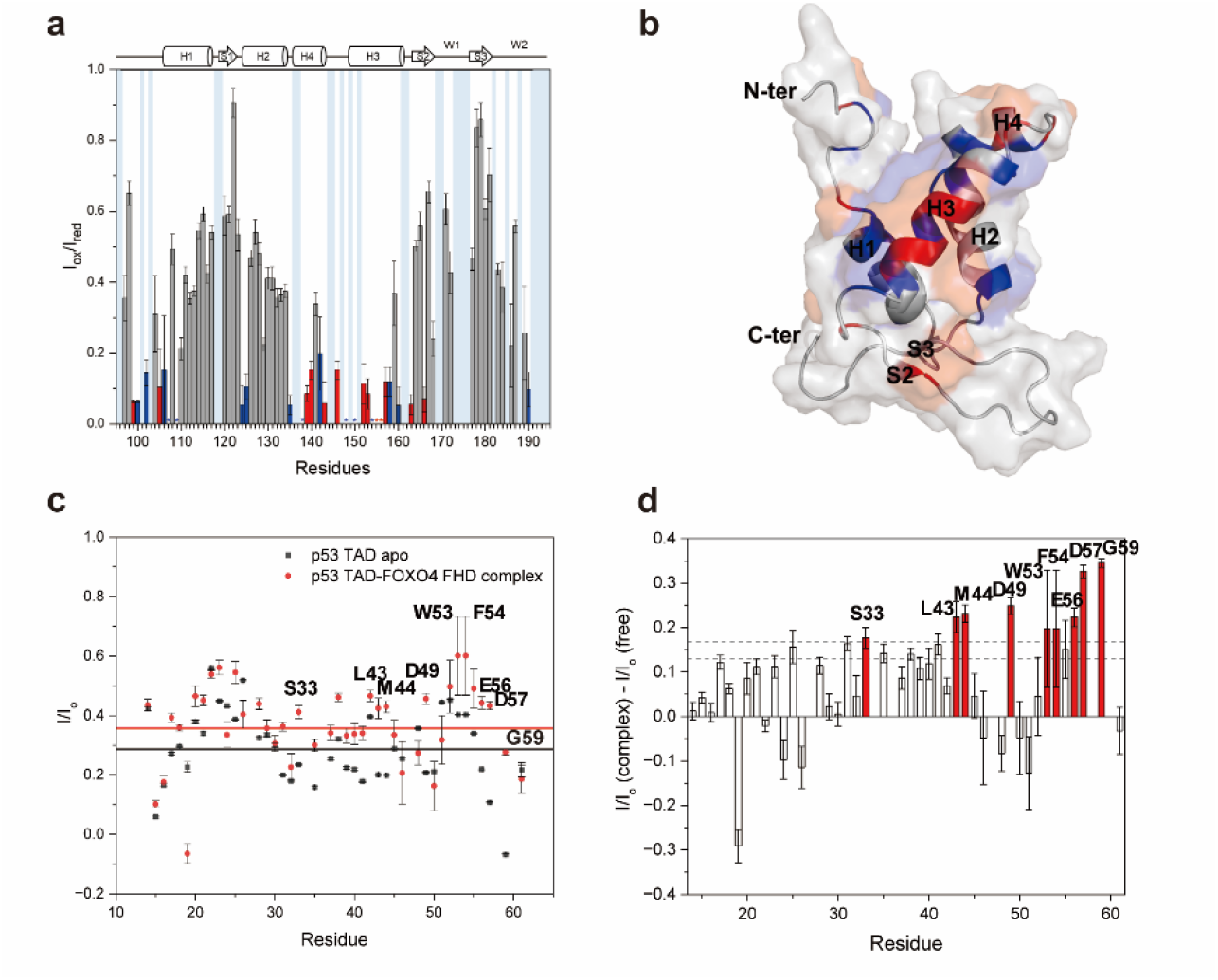
Observation of the FOXO4 FHD and p53 TAD interaction region. Plots of the intensity ratio (I_ox_/I_red_) of FHD signals against the residue number are shown for FHD in complex with TAD modified with MTSL at S46C. (a) I_ox_/I_red_ by residue and their corresponding error bars. Asterisks (*) indicate residues that broadened out under the influence of MTSL. (b) Residues with I_ox_/I_red_ < 0.2 are colored in red, and hydrophobic residues with I_ox_/I_red_ < 0.2 are colored in blue on the NMR structure of FOXO4 FHD (PDB ID: 1E17). (c) HetNOE value of p53 TAD apo and the p53 TAD-FOXO4 FHD complex. (d) Differences in hetNOE values between the p53 TAD-FOXO4 FHD complex and the p53 TAD apo.

A previous CSP analysis using FOXO4 FHD titration of ^15^N-labeled p53 TAD showed that significant perturbations two standard deviations above average were mostly observed in residues within the p53 TAD2 region^38^. This indicates the importance of the p53 TAD2 region in FOXO4 FHD binding. CSP analysis was used to observe the interaction sites based on changes in the surrounding chemical environment. The results imply that even residues not directly involved in the interaction may exhibit changes in their chemical environment if they are adjacent to residues crucial for the interaction or undergo conformational changes upon binding^39,40^. To evaluate the FOXO4 FHD-binding region in p53 TAD using a dynamic tool, we attempted to measure the hetNOE of the apo- and FHD-bound states of p53 TAD. We expected that hetNOE, which provides information on protein flexibility, could identify key residues that directly participate in the interaction by comparing the values of the apo- and FHD-bound states. Initially, a hetNOE experiment was conducted on ^15^N-labeled p53 TAD in the apo state. Typically, a hetNOE value for ^1^H-^15^N the NH bond close to 1 indicates a more rigid conformation of the residue. Regions with secondary structures typically exhibit higher hetNOE values (≥ 0.6). In the p53 TAD apo state, most residues showed low values (< 0.6) (Figure. 2c). This result indicates a disordered protein lacking a secondary structure. The p53 TAD appears to exist in a flexible form without a secondary structure^41^. Subsequently, hetNOE experiments were performed on the complex formed between ^15^N-labeled p53 TAD and unlabeled FOXO4 FHD (1:4). Unlike previous results, the hetNOE values for most residues increased compared to those observed in the apo state of p53 TAD (Figure. 2c). This suggests that p53 TAD and FOXO4 FHD bind together. In particular, an increase in the hetNOE value of > 0.2 was observed for residues corresponding to TAD2 and the C-terminus (Figure. 2d). This result is consistent with those of previous reports, supporting the notion that the TAD2 region plays a substantial role in the interaction with FOXO4 FHD^38^. Conversely, residues within positions 45–52, except for residue 49, did not exhibit a notable increase in the hetNOE value. Residues in the anterior and posterior regions of TAD2 and the C-terminus of p53 TAD appear to be important for the interaction with FOXO4 FHD.

### Hydrophobic residues in p53 TAD2 are critical for interaction with FOXO4 FHD

Based on the significance of both the TAD2 region and the C-terminus of p53 in interacting with the FOXO4 FHD, we attempted to develop an inhibitor targeting the FOXO4-p53 interaction. We designed a peptide inhibitor prototype, named p53 TAD PEP, comprising 16 amino acids corresponding to residues 42–57 of the p53 TAD sequence (Figure. 1). We chose the peptide sequence based on our hetNOE and CSP analyses conducted in previous studies^38^. This sequence spans the region exhibiting a significant perturbation or notable increase in the hetNOE value. To verify the binding capability of p53 TAD PEP to the FOXO4 FHD, we performed FPA. FPA analysis was conducted by increasing the concentration of unlabeled FOXO4 FHD and maintaining a constant concentration of fluorescein isothiocyanate (FITC)-labeled p53 TAD PEP. The results indicate that p53 TAD PEP bound effectively to FOXO4 FHD, with a K_d_ of 0.11 μM (Figure. 3a). Additionally, ^1^H-^15^N heteronuclear single quantum coherence (HSQC) NMR experiments were conducted to verify whether p53 TAD PEP effectively inhibited the interaction between FOXO4 FHD and p53 TAD. We conducted titration experiments on the complex of ^15^N-labeled p53 TAD (13-61) and unlabeled FOXO4 FHD with increasing concentrations of p53 TAD PEP. We observed the dissociation of p53 TAD (13-61) from FOXO4 FHD upon the addition of p53 TAD PEP, resulting in a chemical shift of the ^15^N-p53 TAD-FOXO4 FHD complex towards the unbound state of p53 TAD (Figure. 3b). This finding indicates that p53 TAD PEP effectively competes with p53 TAD for FOXO4 FHD binding. Collectively, these results suggest that p53 TAD PEP functions as a peptide inhibitor that targets the FOXO4-p53 interaction.

**Figure 3.**
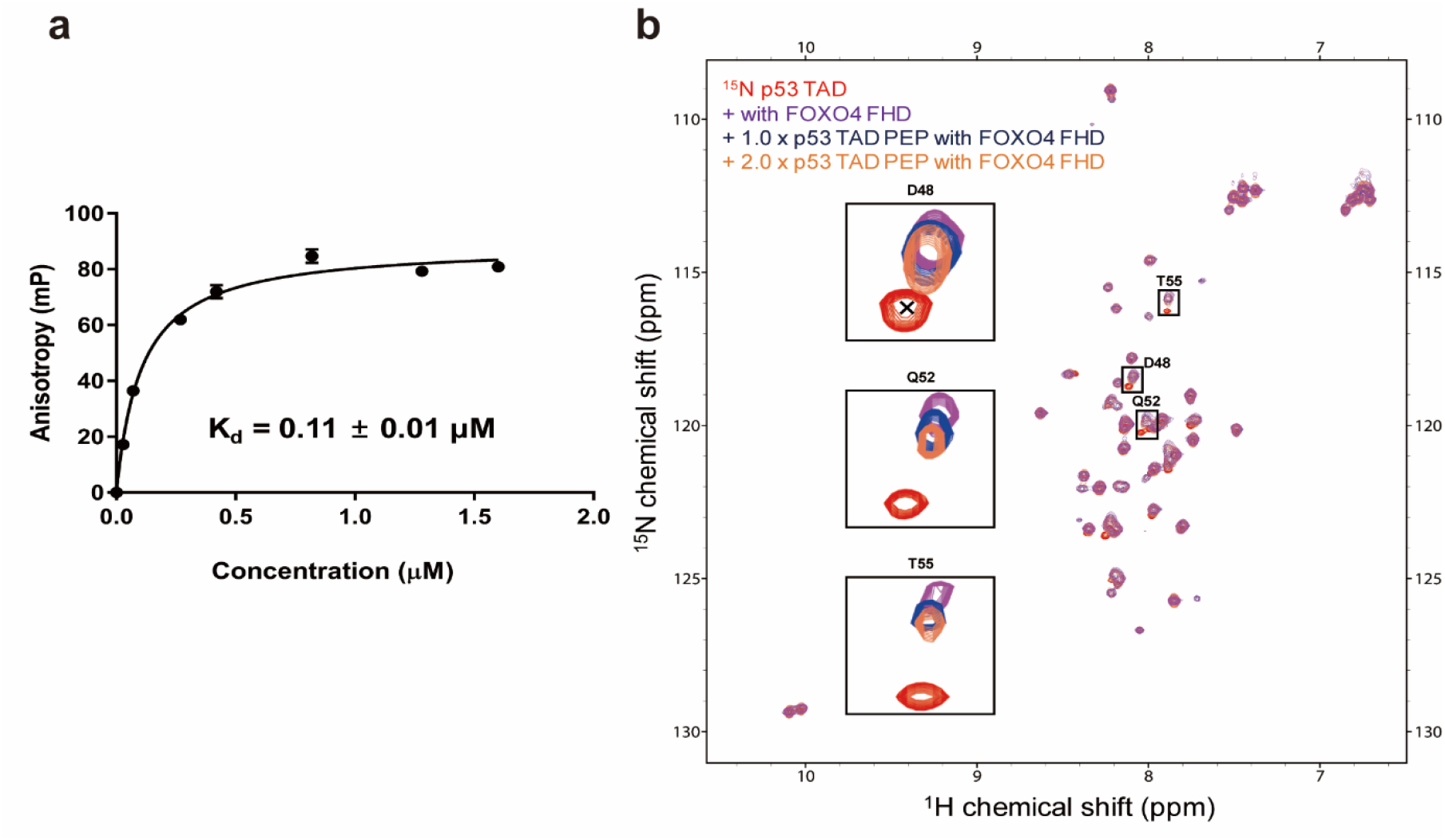
Verification of the inhibitory ability of p53 TAD PEP against the binding of FOXO4 FHD to p53 TAD. (a) FPA experiments were conducted with DNA, measuring the FPA values of FITC-labeled p53 TAD PEP as the concentration of FHD increased. (b) ^1^H-^15^N HSQC NMR spectra were obtained for the ^15^N-labeled FOXO4 FHD-^14^N p53 TAD complex incubated with increasing stoichiometric equivalents of p53 TAD PEP (0, 100 and 200 μM).

To improve the p53 TAD PEP sequence as a competitor, we identified residues essential for FOXO4 FHD binding. Our PRE experiments suggest that hydrophobic interactions play a crucial role in the interaction between FOXO4 FHD and p53 TAD. Based on these observations, we designed peptides in which the individual hydrophobic residues were substituted with alanine. Subsequently, we evaluated the inhibitory ability of the peptide by titrating the concentration of alanine-substituted p53 TAD PEP by combining FITC-labeled p53 TAD PEP and FOXO4 FHD (Figure. S1a). When residues L43, W53, and F54 were substituted with alanine, the inhibitory ability was significantly reduced, indicating that these hydrophobic residues were essential for FOXO4 FHD binding (Table 1). Moreover, a slight reduction in the inhibitory ability was observed when residue M44 was substituted with alanine. This suggests that the hydrophobic residues in p53 TAD significantly contribute to FOXO4 FHD binding.

**Table 1.**
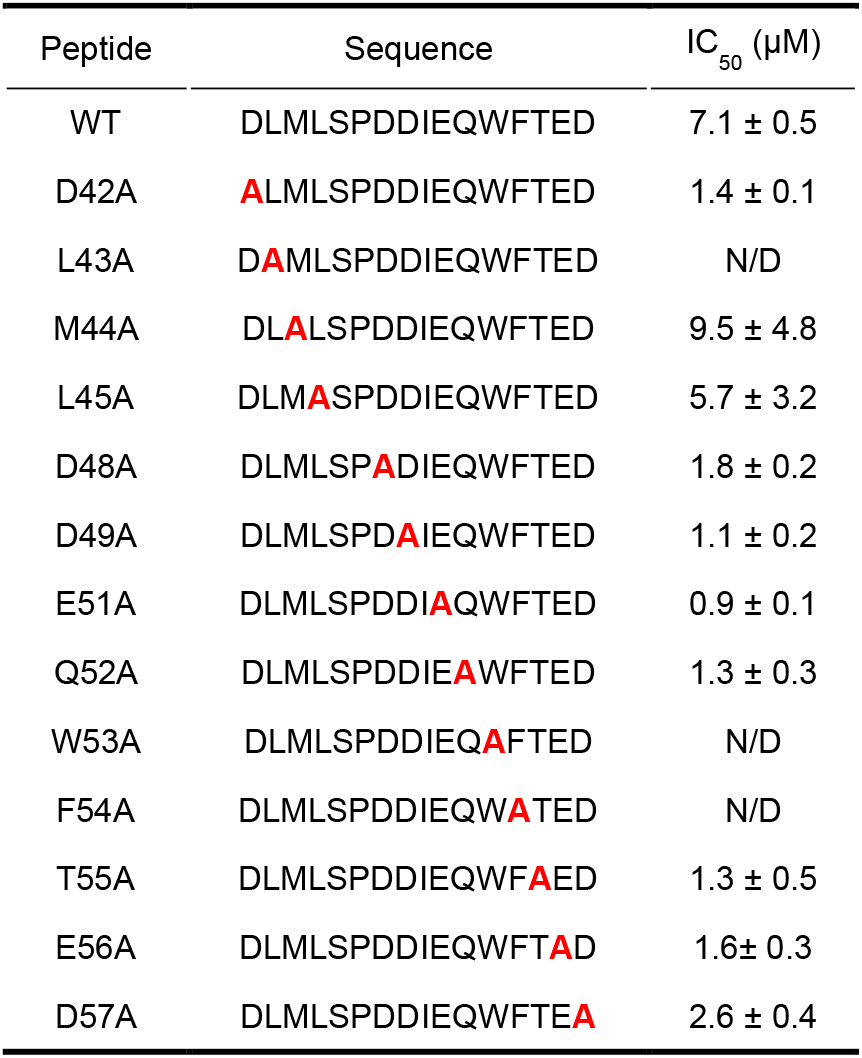
IC_50_ values of wild-type (WT) and alanine-substituted peptides. Alanine mutations are shown in bold red. IC_50_ values were measured using an FPA competition assay.

### A peptide with reduced acidity was designed for increased cell permeability

The p53 TAD2 sequence exhibited a highly negative charge, whereas the p53 binding interface of the FOXO4 FHD contains several positively charged residues. Consequently, it was hypothesized that electrostatic interactions play a significant role in the interactions between these two domains. To test this hypothesis, we designed peptides by substituting negatively charged amino acids and those with a CSP greater than two standard deviations^38^ with alanine. We then evaluated the inhibitory activities of the modified peptides (Figure. S1b, S1c). Interestingly, substituting the amino acids with alanine enhanced the peptide’s inhibitory ability (Table 1). This suggests that these residues are not crucial for FOXO4 FHD binding and may represent candidates for substitution to improve inhibitory ability.

Based on our alanine screening, we aimed to optimize the sequence of p53 TAD PEP to improve its cell permeability. At pH 7.4, the net charge of p53 TAD PEP is -6.2. Since cell membranes are negatively charged, peptide drugs can efficiently penetrate membranes when their overall charge is neutral or positive^42^. The negatively charged residues in the p53 TAD PEP were not crucial for FOXO4 FHD binding (Table 1). Hence, a peptide was designed by substituting the negatively charged residues D42, D48, D49, E51, E56, and D57 within the p53 TAD PEP with alanine to increase its binding affinity to FOXO4 FHD and enhance membrane penetration. The inhibitory ability of the designed peptide (CAND) was then verified using FPA competition analysis to ensure its continued effectiveness in inhibiting the interaction between FOXO4 and p53 TAD (Figure. 4a). Consequently, the inhibitory ability of the modified peptide was enhanced by approximately three-fold compared to that of p53 TAD PEP (Figure. 4b). Thus, we have successfully designed a peptide optimized for cellular applications.

**Figure 4.**
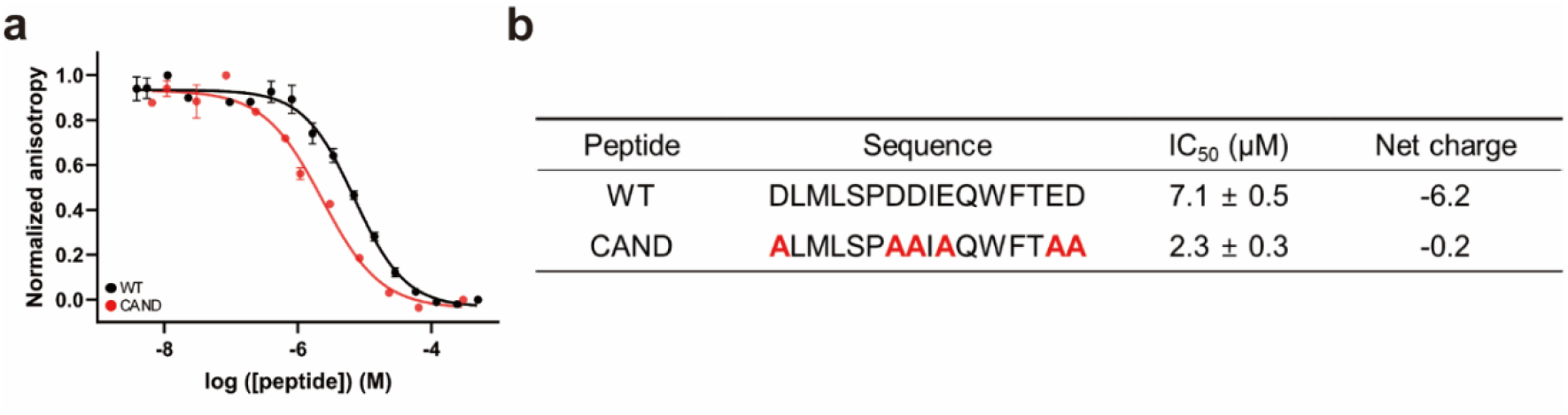
Competitive FPA inhibition assay of peptide candidates with substituted residues. (a) Inhibition of FITC-labeled p53 TAD binding to FOXO4 FHD by increasing concentrations of WT and sequence-optimized peptide candidates (CAND). (b) IC_50_ values and net charges of the WT and CAND peptides at pH 7.4.

### CPP-CAND exhibits selective senolytic activity

Doxorubicin is known to induce senescence in the A375 melanoma cell line^25^. In this study, we induced senescence in A375 cells using doxorubicin and confirmed cellular senescence through senescence-associated beta-galactosidase (SA-β-gal) staining (Figure. 5a). Dividing and senescent cells were treated with the peptides, and the cell viability was measured using the MTS assay, with the IC_50_ values determined accordingly. Both p53 TAD PEP and CAND showed limited efficacy in inducing cell death in both dividing and senescent cells (Figure. S2a, S2b). We then hypothesized that the inefficacy of the peptide in inducing cell death in both types of cells might be attributed to inadequate cellular uptake. To address this, we attempted to enhance the peptide’s cellular uptake. Various techniques, such as incorporating a cell-penetrating peptide (CPP) sequence or lipid conjugation, were employed to facilitate the intracellular delivery of the peptides^43^. Since the net charge of CAND is -0.2, resulting in an overall neutral charge, it is difficult for CAND to be effectively transported to the cell nucleus on its own. Initially, we performed an MTS assay by introducing palmitic acid to CAND (Figure. S2c). However, the peptide with the introduced palmitic acid exhibited a significantly reduced water solubility. Moreover, the peptide demonstrated high toxicity and lacked selectivity. Next, we introduced a CPP sequence to enhance the peptide’s intracellular transport. We designed two peptides to enhance cellular uptake and facilitate nuclear localization by introducing CPP sequences. One peptide contains a hydrophobic sequence at its N-terminus (PFVYLI-CAND)^44^, while the other includes a positively charged, hydrophilic HIV-TAT sequence at its N-terminus (CPP-CAND)^45^ (Figure. 1b, S2g). The PFVYLI-CAND peptide, which includes a hydrophobic CPP sequence, showed no efficacy in inducing cell death in either cell type (Figure. S2d). On the other hand, CPP-CAND, which integrated the positively charged HIV-TAT sequence, demonstrated senolytic activity (Figure. 5b). In our experimental condition, the SI_75_ of CPP-CAND demonstrated significant superiority compared to FOXO4-DRI, with an approximately-1.2 fold higher selectivity, despite its shorter length. The SI_50_ values of CPP-CAND showed comparable efficacy with FOXO4-DRI (Figure. 5c). Treatment of cells with a peptide containing only the HIV-TAT sequence, as well as a peptide where the HIV-TAT sequence was introduced into the N-terminus of p53 TAD PEP, resulted in a minimal induction of cell death (Figure. S2e–S2g). These findings suggest that our peptide exhibits potent functionality within cells as a senolytic drug, with selectivity for senescent cells.

**Figure 5.**
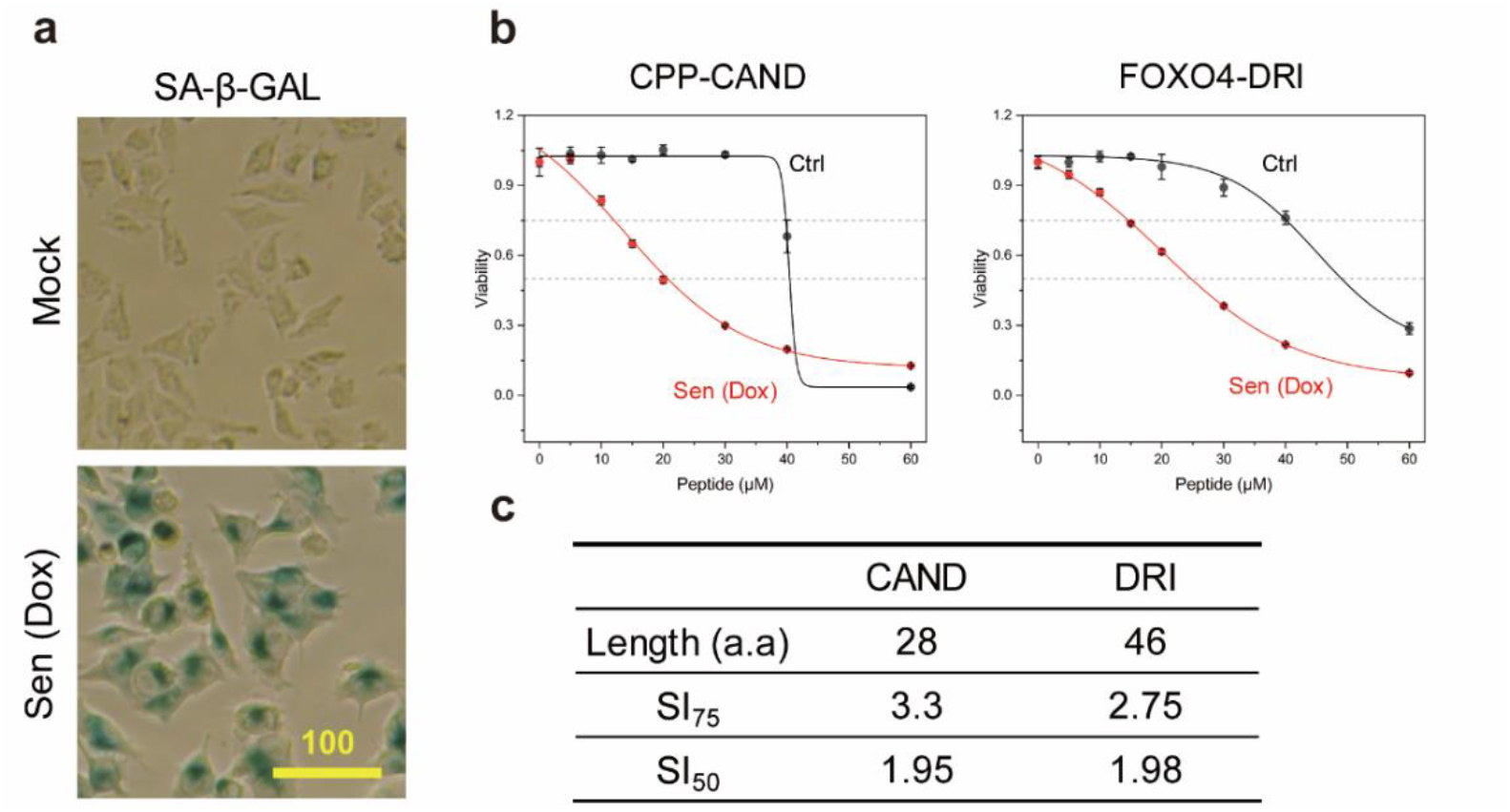
Validation of the candidate peptide in a model of doxorubicin-induced senescence. (a) Images depicting dividing and senescent A375 cells stained for SA-β-gal. (b) Viability assay of senescent and control A375 cells incubated with increasing doses of CPP-CAND and FOXO4-DRI (μM). (c) Selectivity index (SI) of FOXO4-DRI and CPP-CAND. The SI_50_ and SI_75_ values reflect the variations in the IC_50_ and IC_75_ values obtained from non-linear regression analyses for both groups, respectively.

### CPP-CAND disrupts FOXO4-p53 foci and induces apoptosis through caspase-dependent pathways

FOXO4 is progressively recruited to euchromatin foci following senescence induction and is retained in the nucleus of senescent cells through its interaction with p53, contributing to the maintenance of the senescent state^34^. FOXO4-DRI and ES2 peptides effectively disrupted the FOXO4-p53 interaction within the nucleus of senescent cells, leading to selective induction of apoptosis in these cells^25,34^. Since CPP-CAND was designed to interfere with the FOXO4-p53 interaction, we investigated the disruption of FOXO4-p53bp1 foci in the nucleus of senescent cells after CPP-CAND treatment.

Immunofluorescence results showed a significant increase in FOXO4 and p53bp1 foci within the nucleus of senescent cells compared to those in dividing cells (Figure. 6a, 6b). We also observed frequent overlap between FOXO4 and p53bp1 foci, indicating an interaction and co-localization between FOXO4 and p53 in the nuclei of senescent cells (Figure. 6a). This implies that the FOXO4-p53 interaction effectively induces cellular senescence. After treatment with CPP-CAND, a significant reduction of approximately 7.1-fold in the number of FOXO4 foci was observed in the nuclei of the senescent cells. This suggests that CPP-CAND successfully inhibited the FOXO4-p53 interaction in the nucleus (Figure. 6c, 6d). Furthermore, we treated dividing and senescent cells with 20 μM of CPP-CAND and assessed caspase 3/7 activity to examine whether the apoptosis observed was caspase-dependent. Although the same concentration of CPP-CAND was applied to each cell type, caspase 3/7 activity was only observed in 44% of the senescent cells, whereas almost no activity was observed in dividing cells (Figure. 6e). In the control group, in which dividing and senescent cells were treated with DMSO, no caspase 3/7 activity was detected (Figure. S3). Our data clearly showed that caspase-dependent apoptosis was specifically induced by CPP-CAND in senescent cells. Taken together, these results suggest that CPP-CAND inhibited the FOXO4-p53 interaction in the nucleus, which enabled the extranuclear transport of p53 and selectively induced apoptosis in senescent cells.

**Figure 6.**
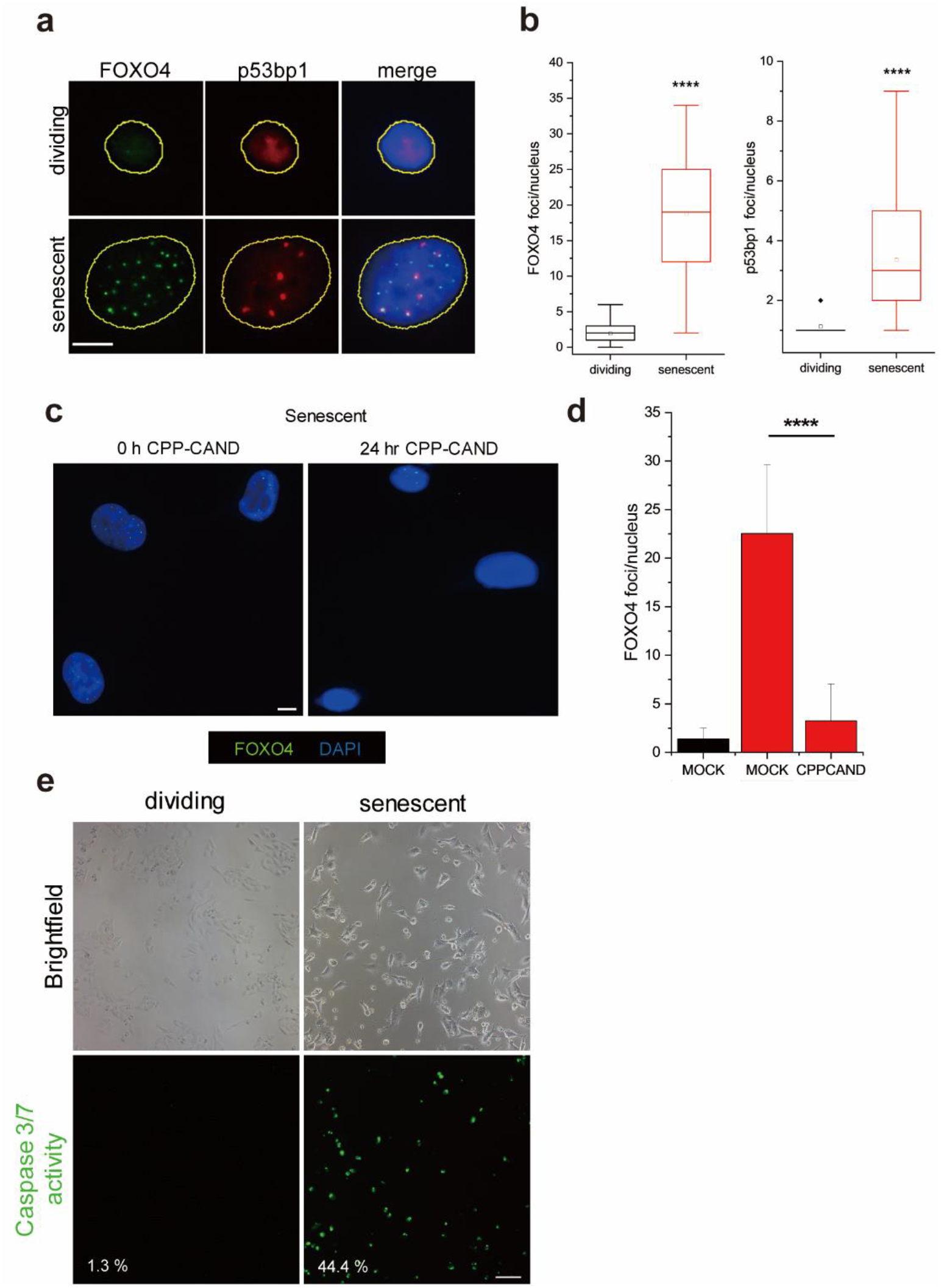
CPP-CAND induces apoptosis by disrupting FOXO4-p53 foci through a caspase-dependent pathway. a. Representative images of FOXO4 and TP53BP1 (p53BP1) foci in dividing and doxorubicin-induced senescent A375 cells. The outline of the nucleus is indicated by the yellow line. b. Box and whisker plots illustrating the distribution of FOXO4 and p53bp1 foci in the nuclei of A375 cells. c. FOXO4 foci in senescent A375 cells treated with 30 μM of CPP-CAND for 24 h. d. Bar graphs of FOXO4 foci/nuclei in A375 cells. Black bars represent dividing cells, while red bars represent senescent cells. The mock group was treated with DMSO as a control. ****p < 0.0001. e. Caspase 3/7 activity was assessed in dividing A375 cells and senescent A375 cells treated with 20 μM CPP-CAND for 24 hours. The percentage of green fluorescent cells per cell is indicated in the panel figures. Scale bar (a,c) = 10 μm, (e) = 100 μm.

Additionally, to verify the efficacy of CPP-CAND in triggering cancer cell senescence induced by other anticancer drugs, we tested its effects on senescent cancer cells induced by cisplatin (CDDP). CDDP induces senescence in A375 cells via the p53/p21 pathway^44^. Initially, senescence was confirmed through SA-β-gal staining following treatment with 4 µM of CDDP (Figure. S4a). Subsequently, both the dividing and senescent cells were treated with CPP-CAND. The SI_75_ value obtained was approximately 2.4, indicating that CPP-CAND also exhibited selectivity for senescent cancer cells induced by other anticancer drugs (Figure. S4b, S4c). These findings suggested that our peptide exhibited potent functionality within cells as a senolytic drug with selectivity for senescent cells.

## Discussion

Cellular senescence, triggered by diverse external and internal stressors, serves as a protective response that prevents the proliferation of impaired cells, thereby conferring resistance to cancer^45,468^. However, the prolonged accumulation of senescent cells can paradoxically induce cancer development and various age-related diseases by inducing an inflammatory environment via the SASP^47-49^. The interaction between FOXO4 and p53 within the nucleus plays a crucial role in the survival of both normal and senescent cancer cells^25,34^. This study employed various experimental methods to develop peptides that inhibit interactions between FOXO4 and p53. We used NMR spectroscopy to identify the crucial region within the p53 TAD responsible for FOXO4 binding (Figure. 2c, 2d). Our findings indicate that hydrophobic interactions contribute more substantially to the interaction between the FOXO4 FHD and p53 TAD than electrostatic I nteractions. Generally, hydrophobic residues form essential contacts during protein-protein interactions, whereas acidic residues mediate long-range interactions with positively charged residues^50-53^. In contrast to these reports, the acidic residues of p53 TAD were not critical for its interaction with the FOXO4 FHD (Table 1). Moreover, neutralizing these charges enhanced the inhibitory activity of the peptides against FOXO4 FHD and p53 TAD. In previous studies, p53 TAD was shown to significantly influence the DNA selectivity of p53 DBD, with the acidic residues of TAD believed to contribute to this selectivity through interactions with the basic residues of DBD^54^. Therefore, the acidic residues of p53 TAD appear to have no significant impact on its interaction with the FOXO4 FHD, as these residues primarily contribute to DNA selectivity rather than binding to the FOXO4 FHD.

Although we found an optimized sequence for the FOXO4 FHD-p53 TAD interaction, it did not show senolytic activity by itself (Figure. S2b). In the present study, cationic CPP was found to be necessary for effective intracellular delivery and senolytic effects (Figure. 5b, S2d). CPPs are peptides characterized by either cationic or hydrophobic motifs that enable them to cross cell membranes, thereby acting as drug carriers. Peptides with cationic motifs utilize electrostatic interactions to target and bind negatively charged phospholipids in cell membranes, thereby facilitating their entry into cells^55^. Peptides with hydrophobic motifs interact well with the lipid components of cell membranes, aiding their ability to penetrate cells^56^. Based on this ability, CPPs effectively deliver various therapeutic agents, including peptide drugs, small-molecule drugs, and nucleic acids, into cells^57^. The requirement for cationic CPP in our study is consistent with findings from other studies, where CPPs with hydrophobic motifs did not significantly enhance cell uptake^60^. This low efficacy may be due to the poor solubility of hydrophobic CPPs^61^. It was assumed that CAND has low solubility owing to the substitution of negatively charged amino acids with alanine. This issue was further aggravated by the introduction of CPPs with hydrophobic motifs, which may explain the ineffectiveness of hydrophobic CPPs for the cellular delivery of CAND. Consequently, cationic CPPs, which are better suited for cellular uptake owing to their positive charge, are more effective for the cellular delivery of CAND.

FOXO4-DRI is a senolytic peptide that targets p53, a crucial tumor suppressor that prevents genomic mutations^34^. Targeting p53 presents a substantial risk of off-target effects. In addition, FOXO4-DRI has a relatively high production cost because of its long length and the requirement for D-type amino acids when synthesized (Table S1). Conversely, CPP-CAND targets FOXO4, which is explicitly expressed in senescent cells, providing a senescent cell-specific target. Despite its relatively short length of 28 amino acids and the requirement for L-type amino acids, it exhibited high selectivity for senescent cells. ES2 is a senolytic peptide that targets FOXO4 CR3 but exhibits relatively poorer selectivity for senescent cells compared to FOXO4-DRI^25^ (Table S1). In addition, CR3 lacks a secondary structure^58,59^. In general, disordered domains are unsuitable targets for potent peptide binding because of the lack of well-defined binding surfaces, conformational flexibility, and associated entropic penalties^60,61^. CPP-CAND targets the well-ordered FOXO4 FHD domain, contributing to its high selectivity for senescent cells. In summary, CPP-CAND is a FOXO4-p53 interaction inhibitor that is anticipated to offer cost-effectiveness, high selectivity, and safety compared to previously developed peptide interaction inhibitors.

A recent proposal suggests employing senolytics as a secondary adjuvant therapy in cancer treatment following a one-two-punch approach^626^. Conventional cancer therapy typically eradicates the majority of cancerous cells while concurrently triggering senescence in the remaining few cells (first punch). Senolytics are then used to eliminate senescent cancer cells (second punch). This strategy holds promise for addressing the vulnerability of conventional cancer therapy resulting from the persistence of senescent cancer cells. In studies employing this approach, the risk of cancer progression and recurrence triggered by senescent cancer cells was significantly mitigated^63^.

CPP-CAND could be applied as a senolytic after confirming its effectiveness across different cell populations and in animal experiments. Future studies should validate the scope and efficacy of CPP-CAND, including its performance in a variety of cancer cell lines and with various senescence inducers. Additionally, *in vivo* experiments should be conducted to assess the drug’s safety, efficacy, long-term effects, and potential side effects. These further studies may enhance the clinical applicability of CPP-CAND and contribute to the treatment of cancer and aging-related diseases. We hope that the peptide drug developed in this study could be a viable option for second-line adjuvant tumor therapy.

## METHOD DETAILS

### Sample preparation

FOXO4 FHD (95-195) was subcloned into the pET His6 Tobacco Etch Virus (TEV) LIC cloning vector (2B-T, obtained from Scott Gradia, Addgene Plasmid #29666) and transformed into BL21 (DE3) cells. p53 TAD (13-61) was subcloned into the pET His 6 glutathione-S-transferase (GST) TEV LIC cloning vector (2G-T, obtained from Scotta Gradia, Addgene plasmid #29707) and transformed into BL21 (DE3) pLysS cells. Cells were grown at 37 °C to an optical density at 600 nm (OD_600_) of 0.7–0.8 in lysogeny broth (LPS solution LB-05) medium, and then 0.5 mM of isopropyl-1-thio-β-D-galactopyranoside (LPS solution IPTG025) was added. The cells were then incubated overnight at 18 °C. Overexpressed proteins were purified on a Ni-NTA column (Cytiva) with an elution buffer containing 50 mM NaH_2_PO_4_, 300 mM NaCl, and 300 mM imidazole. His6- and GST-tagged protein samples were cleaved using TEV protease, followed by repeated purification on an Ni-NTA column. To further purify FOXO4 FHD and p53 TAD, gel filtration chromatography was conducted using Hi-Load 16/600 superdex 200pg and 75pg (Cytiva) on AKTA pure and AKTA prime systems (Cytiva). The columns were pre-equilibrated with a buffer containing 20 mM HEPES (pH 7.0) and 150 mM NaCl. For NMR studies, the cells were grown in M9 medium with ^15^NH_4_Cl to yield ^15^N-labeled proteins. All peptides, including fluorescein isothiocyanate (FITC)-labeled p53 TAD (42-57), were manufactured by Dandicure (Ochang, Korea) at a purity of 95%.

### NMR spectroscopy

NMR experiments were performed using a Bruker 900 MHz NMR spectrometer equipped with a cryogenic probe (KBSI, Ochang) and a Bruker 600 MHz NMR spectrometer equipped with a cryogenic probe (GIST, Gwangju). All experiments were performed at 25 °C in a buffer solution containing 20 mM HEPES (pH 7.0), 150 mM NaCl, and 1 mM DTT. The amide nitrogen and protons of FOXO4 FHD and p53 TAD have been assigned previously^64,65^. NMR experimental data were processed using Topspin (Bruker) and analyzed using the POKY script^66^.

^1^H-^15^N heteronuclear Overhauser effect (hetNOE) measurements were recorded using a Bruker 900 MHz NMR spectrometer with a cryogenic probe (KBSI, Ochang). The saturation period for protons (^1^H) was set at 5 s. The hetNOE values were derived by comparing the peak heights of the spectral sets.

Paramagnetic relaxation enhancement (PRE) experiments were performed using a Bruker 600 MHz NMR spectrometer with a cryogenic probe (GIST, Gwangju). The paramagnetic probe, S-(1-oxyl-2,2,5,5-tetramethyl-2,5-dihydro-1H-pyrrol-3-yl)methyl methanesulfonothioate (MTSL), was used to label the cysteine residue 46 of p53 TAD^67^, where 300 µM MTSL-labeled p53 TAD was added to 150 µM ^15^N-labeled FOXO4 FHD. The experiments were performed in a buffer solution containing 20 mM HEPES (pH 7.0) and 150 mM NaCl. The peak intensities were measured and plotted as the ratio of the intensities in the oxidized state (I_ox_, paramagnetic, without ascorbate) to those in the reduced state (I_red_, diamagnetic, with ascorbate). Standard errors for the I_ox_/I_red_ values were calculated by propagating the signal-to-noise ratio of the individual spectra using the equation below^68^:

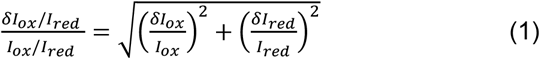

where δI_ox_/I_red_ is the calculated error in the I_ox_/I_red_ ratio, δI_ox_ is the noise level of the oxidized spectrum, and δI_red_ is the noise level of the reduced spectrum. For ^1^H-^15^N heteronuclear single quantum coherence (HSQC) titration experiments, the concentration of p53 TAD PEP was increased while maintaining the same ratio of ^14^N-FOXO4 FHD and ^15^N labeled-p53 TAD.

### Fluorescence polarization anisotropy assay

Fluorescence polarization assays were conducted using a Cytation 5 cell imaging multi-mode reader (BioTek) equipped with a green FP filter at 25 °C, utilizing 96-well microplates (SPL 30496). Each assay was repeated three times and the results were averaged. All experiments were conducted in a buffer solution containing 20 mM HEPES (pH 7.0), 150 mM NaCl, 1 mM dithiothreitol (DTT), and 5% dimethyl sulfoxide (DMSO). To determine the dissociation constant (K_d_) of FHD with p53 TAD PEP, we mixed 10 nM FITC-labeled p53 TAD PEP with increasing concentrations of FHD. The plates were equilibrated for 2 h at 25 °C. The FPA value was determined by measuring the absorbance at an excitation wavelength of 485 nm and emission wavelength of 528 nm and calculated according to the following formula:

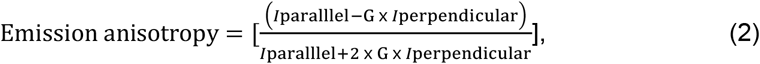

In competitive inhibition experiments, increasing concentrations of inhibitor peptides were titrated with a mixture of 120 nM FHD and 10 nM FITC-p53 TAD PEP. The IC_50_ value, which represents half the maximum concentration of a substance that inhibits a specific biological function *in vitro*, was determined using this assay.

### Cell culture

Human malignant melanoma (A375) cells were obtained from the American Type Culture Collection (ATCC, CRL-1619IG-2) and used within ten passages for all experiments. Cells were cultured in Dulbecco’s Modified Eagle’s medium (Welgene LM001-05) supplemented with 10% fetal bovine serum (Gibco 16000-044), and 1% penicillin/streptomycin (Gibco 15140-122). Cells were incubated in a humidity-controlled incubator at 37 °C with 5% CO_2_. For the detachment of adherent cells, TrypLE™ Express Enzyme (1×) with Phenol Red (Gibco 12605010) was used. Cell lines were regularly tested for mycoplasma contamination using a MycoStrip™ Mycoplasma Detection Kit (InvivoGen rep-mys-10).

Senescent cells were established via DOX-induced senescence. A375 cells were cultured until 30–50% confluence, then treated with a complete growth medium containing 0.1 μM doxorubicin (Sigma D1515). After a 2-day treatment period, the cells were allowed to recover in fresh growth medium for 4 d before further experimentation. A senescence β-galactosidase assay was conducted using the senescence β-galactosidase staining kit (Cell Signaling Technology, 9860) according to the manufacturer’s instructions, and then stained cells were imaged using a ZEISS Primovert inverted microscope (Carl Zeiss Microscopy).

### Cell viability measurements

For this experiment, approximately 12,000 senescent and 6,000 non-senescent cells were seeded in triplicate wells of 96-well plates. After one day of adhesion, the plates were treated with various concentrations of the peptide, with daily replenishment over a total of two days. Cell proliferation was evaluated using a Promega CellTiter 96™ AQueous One Solution Cell Proliferation Assay (MTS) as per the manufacturer’s instructions (Promega G3580). The absorbance (490 nm) was measured three hours after incubation using a microplate reader (Tecan).

### Immunofluorescence assays

For all immunofluorescence assays, cells (typically 15,000) were seeded per well of 8-well plates (SPL 30408). Following each experiment, the cells were washed once with PBS (LPS solution CBP007B) and fixed with 10% formalin solution (Sigma HT501128-4L) for 10 min at room temperature. After fixation, the cells were washed with PBS and permeabilized with 0.1% Triton X-100 (Daejung 8566-4405) in PBS for 10 min at room temperature. Subsequently, the cells were washed with PBS and blocked for 1 h with 1% bovine serum albumin (BSA, Sigma A9418) and 22.52 mg/mL glycine in PBST (PBS + 0.05% Tween-20, LPS Solution CBP007T). Primary antibodies against FOXO4 (Cell Signaling Technology 9472) and TP53BP1 (BD Biosciences 612523) were diluted in 1% BSA in PBST and incubated with the cells overnight at 4°C. After incubation with the primary antibody, the cells were washed with PBST and stained with secondary antibodies (Goat Anti-Rabbit IgG H&L Alexa Fluor 488, Abcam, ab150077; Goat Anti-Mouse IgG H&L Alexa Fluor 594, Abcam, ab150116) diluted in 1% BSA in PBST for 1 h at room temperature. After incubation with the secondary antibody, the cells were washed with PBST and mounted using a Fluoroshield Mounting Medium with DAPI (Abcam ab104139). For experiments involving caspase-3/7 staining, CellEvent Caspase-3/7 Green Detection Reagent (Thermo Scientific C10723) and CPP-CAND were added to live cells 24 h before imaging using an Eclipse Ti-U inverted microscope (NIKON).

## Acknowledgments

We thank the high-field NMR facility at the Korea Basic Science Institute (KBSI, Ochang) and the GIST Advanced Institute of Instrumental Analysis (GIST, Gwangju) for allowing us to use their NMR spectrometers. We thank Dr. Hyejin Ham (GIST Department of Chemistry) for technical assistance with the cellular experiments. This work was supported by the Yuhan Innovation Program and the National Research Foundation of Korea (NRF) [2021R1A2C1004669 and RS-2024-00411137], funded by the Korean government (MSIT) and by the KBSI under the R&D program [Project No. A412550], supervised by the Ministry of Science and ICT, Korea.

## Competing Interests

The authors declare the following competing interests: [Donghoon Kang and Chin-Ju Park] are listed as an inventor on a pending patent application related to ‘A peptide inhibitor for senescent cell elimination, and a use thereof,’ filed by [Gwangju institute of science and technology] under application number KR10-2024-0086615.

## Data availability

The protein structure from accession codes 1e17 (FOXO4 FHD) was obtained from the PDB at https://www.rcsb.org/. All other data are available from the corresponding author upon reasonable request.

## Author Contributions

D.K. – Conceptualization, Formal Analysis, Investigation, Visualization, Writing-original draft, Writing-review & editing

Y.L. – Formal Analysis, Investigation, Validation

D.A. – Formal Analysis, Investigation, Validation

J.L. – Formal Analysis, Validation

C.-J.P. – Conceptualization, Funding acquisition, Supervision, Writing-review & editing

## Notes

### Competing Interest Statement

The authors have declared no competing interest.

