## Supplementary_data for "Peptide inhibitors targeting FOXO4-p53 interactions and inducing senescent cancer cell-specific apoptosis"

Supplementary Information for “Development of peptide inhibitors targeting FOXO4-p53 interactions and inducing senescent cancer cell-specific apoptosis”

Donghoon Kang ^a^, Yeji Lim ^a^, Dabin Ahn ^a^, Jaeseok Lee ^a^ and Chin-Ju Park ^a*^

a	Department of Chemistry, Gwangju Institute of Science and Technology, Gwangju, 61005, Korea

**^*^ Correspondence:**
Chin-Ju Park

**Supplementary Information**

Results	1

Figure S1. Competitive FPA inhibition assay of alanine-substituted peptides	1

Figure S2. Validation of candidate peptides in a doxorubicin-induced senescence model……….. 2

Figure S3. Validation of caspase 3/7 activity induction using DMSO as control	3

Figure S4. Validation of candidate peptides in a CDDP-induced senescence model……………... 4

**Results**

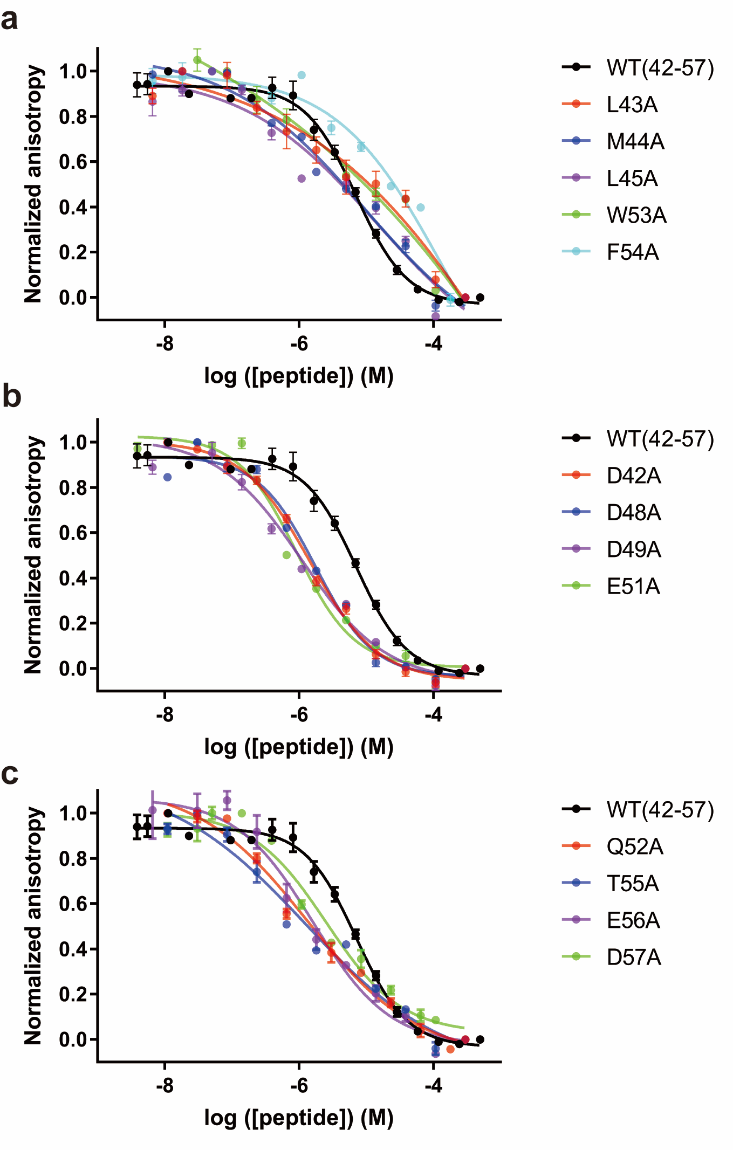
**Figure S1. Competitive FPA inhibition assay of alanine-substituted peptides.**

FITC-labeled p53 TAD binding to FOXO4 FHD was inhibited by increasing the concentration of alanine-substituted peptides.

**Figure S2. Validation of candidate peptides in a DOX-induced senescence model.**

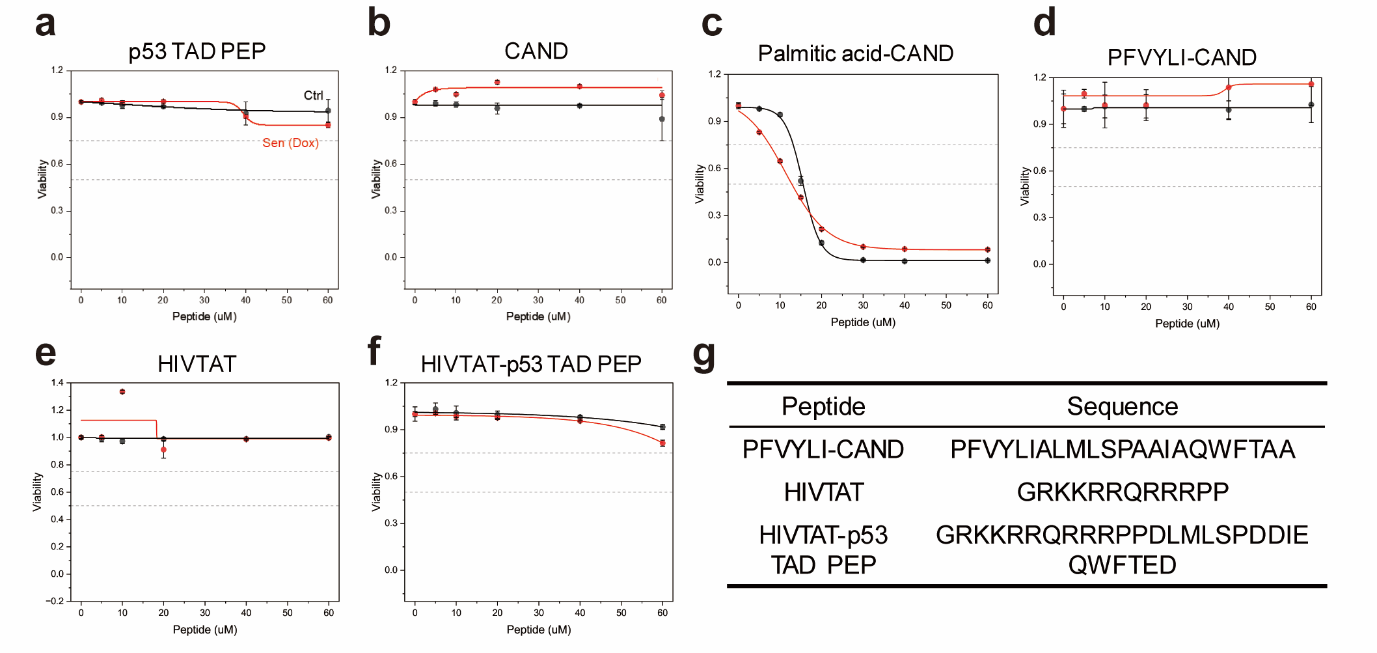

a-f. Viability assay of senescent and control A375 cells incubated with increasing peptide doses (μM). g. Sequences of PFVYLI-CAND, HIVTAT, HIVTAT-p53 TAD PEP peptides.

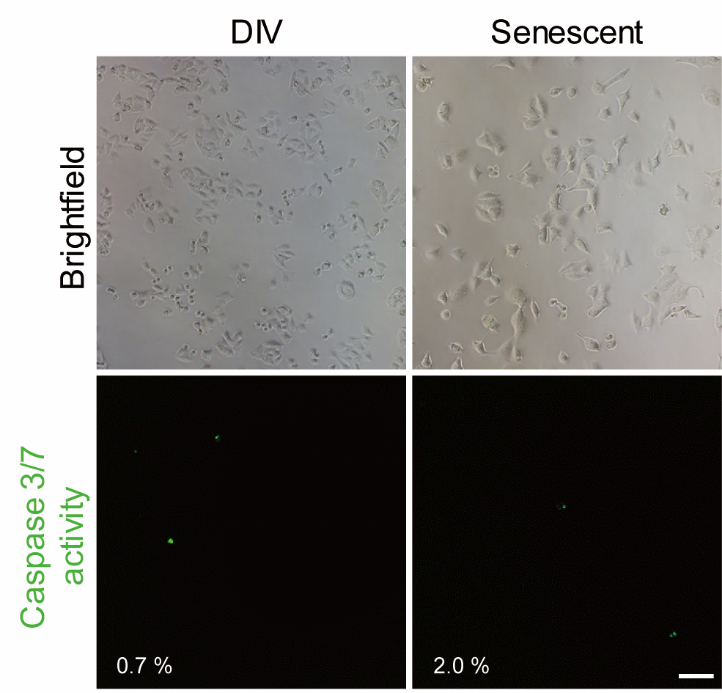
**Figure S3. Validation of caspase 3/7 activity induction using DMSO as a control.**

Caspase 3/7 activity was assessed in dividing and senescent A375 cells treated with DMSO for 24 h. The percentage of green fluorescent cells per cell is shown in the panels. Scale bar = 100 μm.

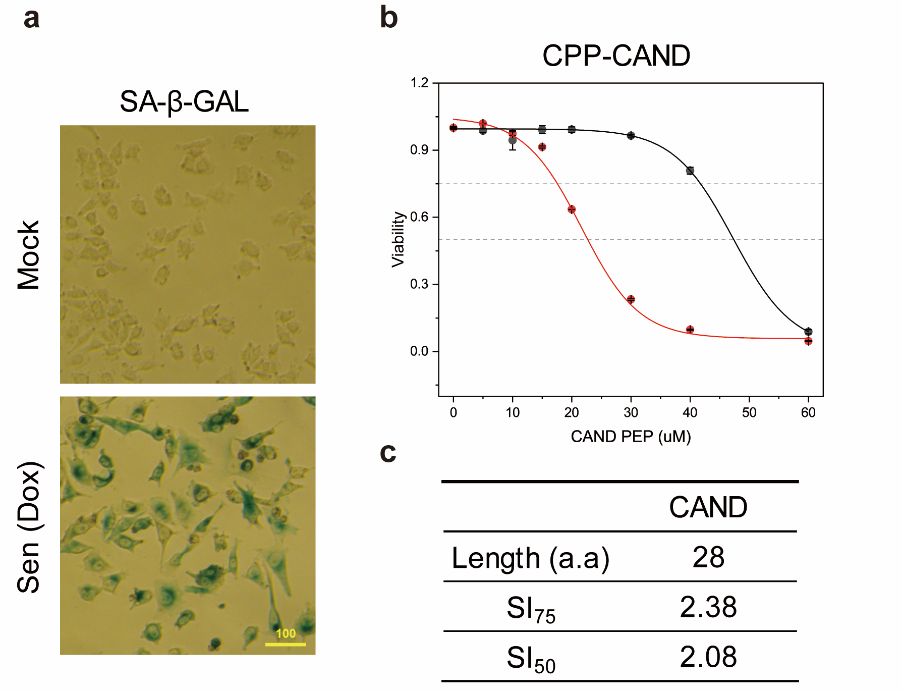
**Figure S4. Validation of candidate peptides in a CDDP-induced senescence model.**

a. Images depicting dividing and senescent A375 cells stained for SA-β-gal. b. Viability assay of senescent and control A375 cells incubated with increasing doses of CPP-CAND and FOXO4-DRI (μM). c. Selectivity index (SI) of FOXO4-DRI and CPP-CAND. SI_50_ and SI_75_ reflect the variations in IC_50_ and IC_75_ values obtained from nonlinear regression analyses for both groups, respectively.
